# A Novel Computational Pre-Procedural Planning Model for Coronary Interventions Based on Coronary CT Angiography

**DOI:** 10.1101/2024.07.29.605713

**Authors:** Mengzhe Lyu, Ce Liang, Xuehuan Zhang, Xiao Wang, Qiaoqiao Li, Ryo Torii, Yiannis Ventikos, Duanduan Chen

## Abstract

In percutaneous coronary intervention (PCI), the ability to predict post-PCI fractional flow reserve (FFR) and stented vessel informs procedural planning. However, highly precise and effective methods to quantitatively simulate coronary intervention are lacking. This study developed a validated virtual coronary intervention (VCI) technique for non-invasive physiological and anatomical assessment of PCI. In this study, patients with substantial lesions (pre-PCI FFR of less than 0.80) were enrolled. VCI framework was used to predict vessel reshape and post-PCI FFR. The accuracy of predicted post-VCI FFR, luminal cross-sectional area (CSA) and centreline curvature was validated with post-PCI computed tomography (CT) angiography datasets. Overall, 21 patients were selected for the study, of which 9 patients (9 vessels) were included in the analysis. The average time for PCI simulation was 24.92 ± 1.00 s on a single processor. The calculated post-PCI FFR was 0.92 ± 0.09 and the predicted post-VCI FFR was 0.90 ± 0.08 (mean difference: -0.02 ± 0.05 FFR unit; limits of agreement: -0.08 to 0.05). Morphologically, the predicted CSA is 16.36 ± 4.41 mm^2^ and post-CSA is 17.91 ± 4.84 mm^2^ (mean difference: -1.55 ± 1.89 mm^2^; limits of agreement: -5.22 to 2.12), the predicted centreline curvature of stented region is 0.15 ± 0.04 mm□^1^ and post-PCI centreline curvature is 0.17 ± 0.03 mm□^1^ (mean difference: -0.02 ± 0.06 mm□^1^; limits of agreement: -0.12 to 0.09). The proposed VCI technique achieves non-invasive pre-procedural anatomical and physiological assessment of coronary intervention. The proposed model has the potential to optimize PCI pre-procedural planning and improve the safety and efficiency of PCI.

**Highlights:**

- Present a computational pre-procedural planning model for coronary interventions.
- Develop a computational framework to predict post-PCI FFR.
- Validation of the model with post-PCI CT angiography datasets.
- The proposed model has the potential to optimize PCI pre-procedural planning.

## 1. Introduction

The observation of fractional flow reserve (FFR) instantly after stent deployment can be used to evaluate functional revascularization gained from percutaneous coronary intervention (PCI) [1]. A better prognosis has been demonstrated for patients with high FFR readings following PCI than for those with low post-PCI FFR [2–4]. Patients with a post-treatment FFR of less than 0.90 may be more likely to develop major adverse cardiac events (MACE) at follow-up, and a higher post-PCI FFR is related to better PCI outcome [5–7]. It has been previously observed that after PCI, approximately one-third of patients who still have inadequate FFR and angina persists for 20% to 50% of patients [8]. Additionally, complications of PCI may affect patient survival and healthcare costs [9–11]. There is a positive correlation between patient risk and the procedural complexity of PCI. It is crucial to be aware of the potential complications of PCI, such as hemodynamic collapse, entrapped equipment, dissections, etc. [12]. As a result, the development of intracoronary physiological and accurate stent deployment assessment tools that help to determine functional revascularization and vessel remodelling have the potential to improve the efficiency and safety of PCI treatment.

Coronary computed tomography (CT) angiography-based fractional flow reserve (FFR_CT_) is a non-invasive physiologic simulation technique that models blood flow based on computed tomography angiogram (CTA) images. As for coronary stenting, research studies on computational patient-specific coronary stenting have received a significant investment recently [13]. The finite element (FE) method is typically used in coronary structural investigations and coronary artery bifurcation stenting [13–15]. Prior studies have made substantial contributions to PCI treatment planning, but they are limited in offering clinical practice assistance in real time. Advanced computational techniques that are able to accurately mimic the morphological and hemodynamic changes in the coronary artery caused by the stent in real time are required. One of the solutions is to use virtual coronary intervention (VCI) tools, which aim to give time-efficient simulations for treatment plans [16]. Pioneered by Heartflow Inc., Redwood City, California, FFR_CT_ Planner is a novel tool that can predict post-PCI FFR in high accuracy [1]. Based on the angiogram, Gosling *et al*. [16] developed a treatment planning tool which can accurately predict virtual (vFFR) with a reasonable computational time.

While these studies make significant contributions to PCI planning, the virtual stenting process is implemented by employing a cubic spline to adjust the cross-sectional radius. This adjustment aims to smooth the vessel trajectory based on the dimensions (diameter and length) of the stent. Consequently, the vessel undergoes modifications, and post-PCI FFR is calculated based on the newly created vessel model. There are some limitations in the existing fast coronary virtual stenting method, as there are not physical-oriented models and the balloon is not included in the simulation process. The balloon is frequently used as a pre-dilation, and post-dilation device during the PCI process and it is used to expand the stent. Besides, vessel-stent interaction can affect the simulation results depending on various situations. To develop a more realistic VCI framework, a balloon should be included, and an efficient contact model should be developed along with the dynamic mesh [17] to accurately simulate the balloon expansion process and balloon-attached coronary reshaping with rapid computation time. Therefore, the overall aim of this study is to develop a fast VCI technique that provides intracoronary post-stenting physiological assessment along with anatomical assessment.

## 2. Methods

### 2.1. Study design

The patients who presented with mild-to-moderate chest pain at the Cardiology Department of Chinese PLA General Hospital from January 2021 to March 2021 were included in this study. Twenty-one patients were initially selected randomly, who had undergone a coronary CTA before and after PCI. The computational fluid dynamics (CFD) procedure was applied to the segmented coronary model to derive pre-PCI FFR and post-PCI FFR. Patients with pre-PCI FFR greater than 0.8 (indicating non-severe stenosis), who experienced failed automated image segmentation, were rejected for post-FFR analysis due to issues primarily related to poor vessel segmentation (such as vessel disconnection), as illustrated in Supplemental Figure 1.

9 patients with substantial lesions determined by pre-PCI FFR of less than 0.80 were considered for further study. The patient characteristics and vessel-specific characteristics are shown in Supplemental Table 1. This study was approved by the Institutional Review Board of the Chinese PLA General Hospital (S201703601). The virtual balloon and stent expansion process is integrated with flow analysis to form the VCI framework, a novel tool that models the PCI process and predicts post-FFR with the change in vessel lumen, as illustrated in Fig. 1. This computational method includes 5 steps, aiming to achieve both non-invasive physiological assessment and detailed anatomical assessment for PCI. The VCI framework has the potential to provide risk evaluation for PCI procedures.

**Figure 1.**
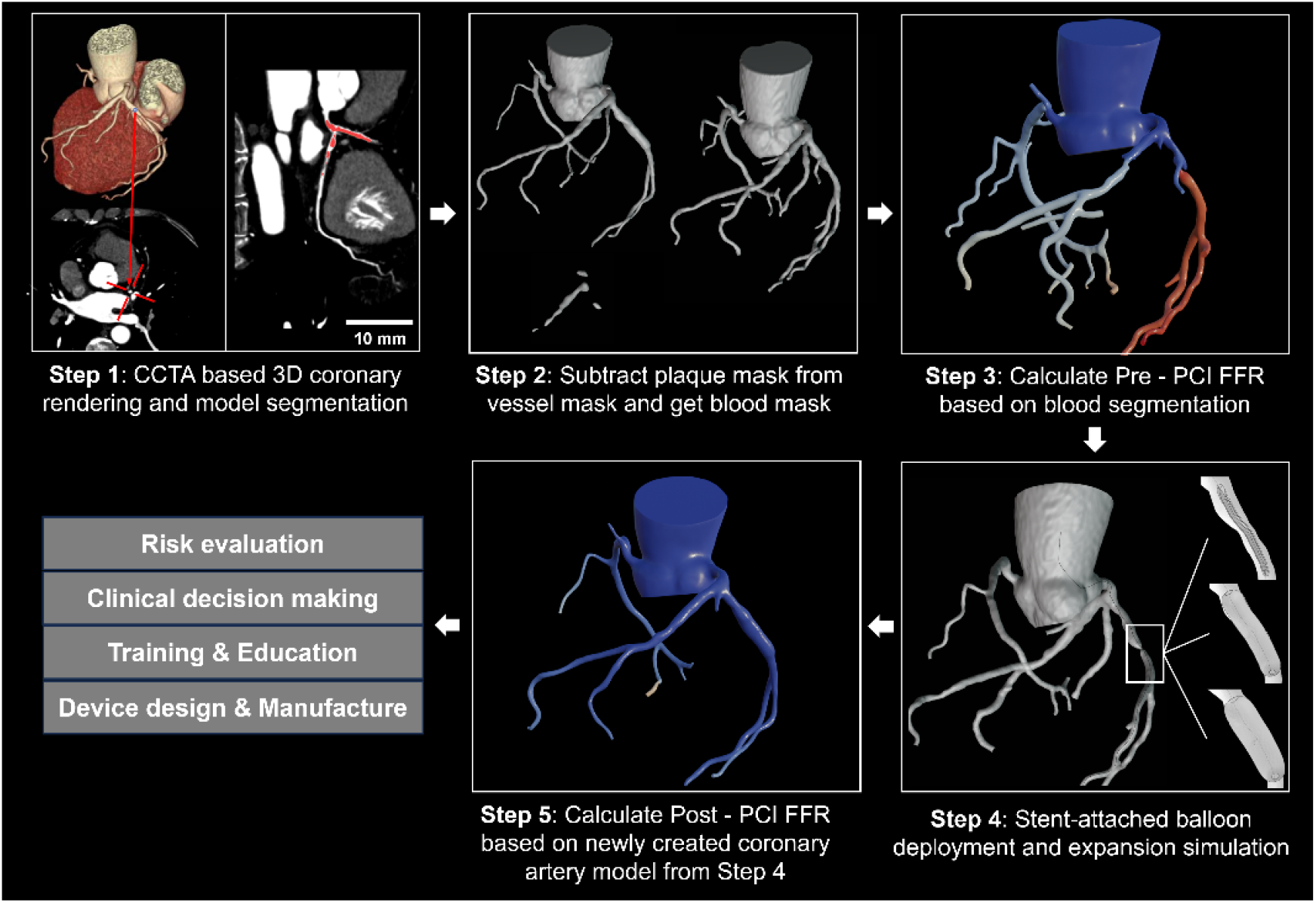
Workflow of patient-specific computational virtual coronary intervention (VCI) framework, overall steps required for the calculation of post-PCI fractional flow reserve (FFR) and risk evaluations. The coronary artery model and plaque were 3D reconstructed and then used to create blood mask (Step 1 and 2) based on coronary computed tomography angiogram (CCTA). The blood masks are meshed and assigned with patient-specific boundary conditions based on CCTA imaging (Step 3). The computationally created balloon and stent are positioned in the region of vessel lesion (Step 4). Based on dynamic mesh, the stenting process is performed computationally. The newly created coronary artery model after stenting process is meshed and assigned with patient-specific boundary conditions again for the calculation of Post-PCI FFR (Step 5).

### 2.2. Coronary CTA image acquisition, automated geometry reconstruction and post-processing

CTA datasets before and after PCI were obtained using a dual-source CT scanner (SOMATOM Definition Flash, Siemens, Germany). The images were acquired with an injection of 70-90 ml of contrast and 50 ml of saline chaser; a threshold of 80 HU; rotation speed of 500 ms; collimation of 64; slice thickness of 1.0 mm; pitch of 1.0; voltage of 100 kV; and current of 200–350 mA. Automated image segmentation was provided by Shukun (Beijing) Network Technology Co. Ltd [18]. The 3D reconstruction of the coronary artery tree model can be seen in Fig. 2(a). The subtraction of the vessel mask and plaque mask gives the blood mask, which is used for surface reconstruction. Surface reconstruction was achieved using an in-house algorithm and saved as STL format files. The balloons were created computationally inside the VCI framework in their crimped state, and the stents used were created based on dimensions provided by manufacturers in their nominal dimensions, as can be seen in Fig. 2(b) and (e). Supplemental Figure 2 shows the overview of virtual fast pre-operative processes.

**Figure 2.**
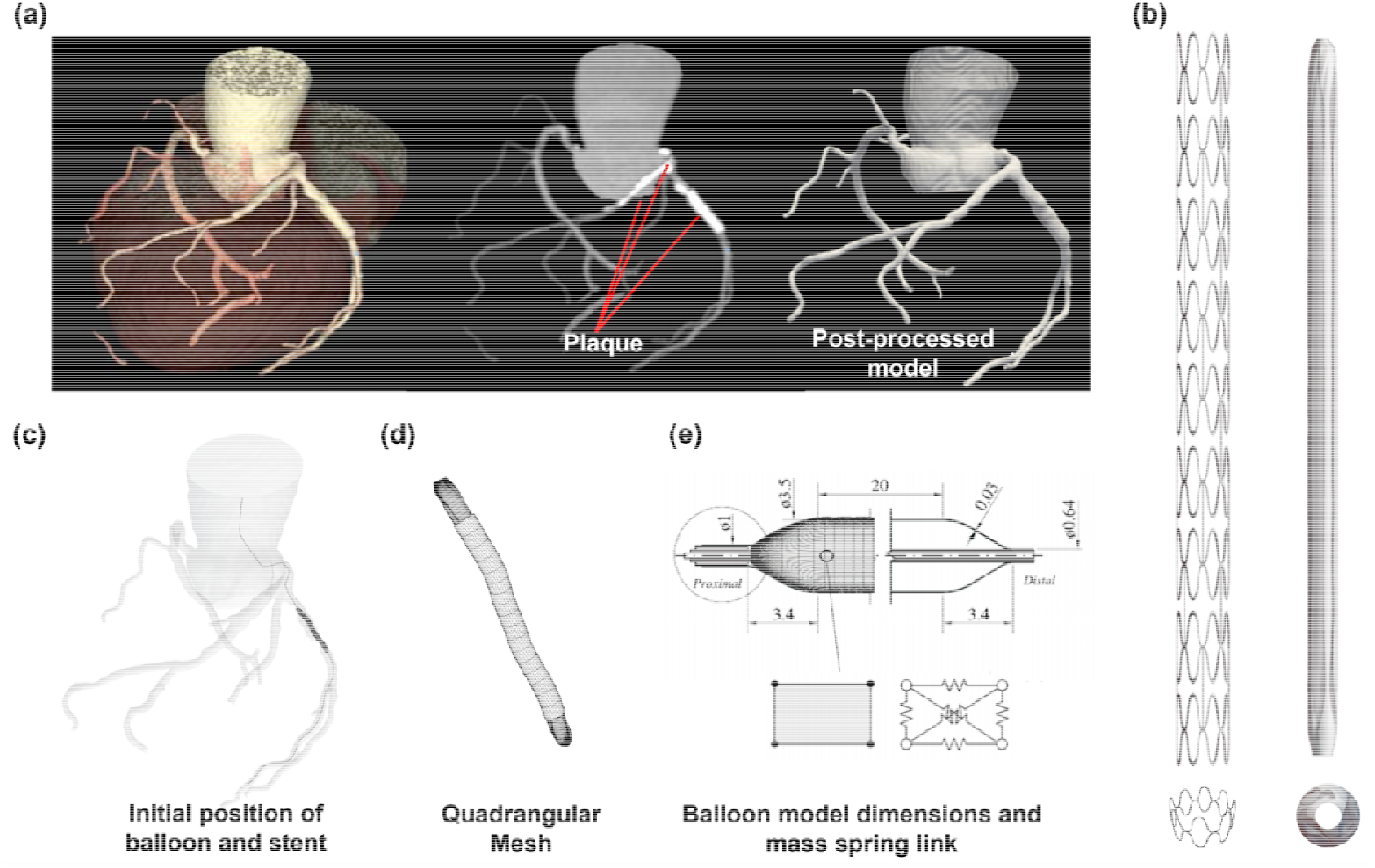
Models (coronary artery, balloon and stent) preparation for VCI framework. (a) 3D automated reconstruction of coronary and plaques based on CCTA dataset, the substruction of plaque mask from vessel mask gives blood mask (post-processed model) which is used for further simulations. (b) Balloon and stent computationally created. (c) Initial position of balloon and crimped stent surface. (d) Quadrangular mesh of balloon and stent. (e) The dimensions of balloon and mass-link used to reconstruct the models.

### 2.3. VCI technique

#### 2.3.1. Theoretical framework

The VCI technique based on the dynamic mesh is used to model the vessel-stent and stent-vessel interactions. The concept of the dynamic mesh, first introduced in the 1990s [19], is considered the earliest and most classic physical model to have been proposed and applied. In the VCI technique, the models are discretized into particle systems that are linked with fictitious mass, damping, and stiffness elements [20]. The force acting on the spring element obeys Hooke’s law. The movement of this dynamic mesh obeys the motion equation of dynamic equilibrium, as shown in the equation below:

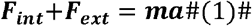

Where *F*_*int*_ is the internal force and *F*_*ext*_ is the external force, m is the mass of each vertex *V*_*i*_, a is the acceleration caused by forces *F*_*int*_ and *F*_*ext*_. The internal force is the resultant force caused by spring pressure of the balloon and damping force (*F*_*Damping*_= −*Dv*), where D is the system damping and damper. The external force corresponds to the contact force of different models, the internal coefficient and it is set to 0.9. Such a damping force and damping coefficient (0.9) are frequently used in virtual stenting applications (such as in the work by [21,22]).

The Euler method can be applied to compute the force applied to the vertex *V*_*i*_ at any given time t. The fundamental equation of dynamics can therefore be solved using an implicit scheme through time. Objects such as the balloon, vessel, and stent are modeled as a mass-spring-damper system, where all the mass nodes in the model are linked with massless springs. Given the diagonal (lumped) mass-matrix *M* ∈ *R*^3*m*×3*m*^, implicit Euler time integration results in the following update rules:

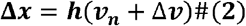

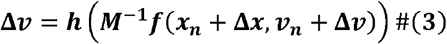

where h is the time interval. The force is evaluated at the end of the time step, known as Backward Euler. Equations 4 and 5 are equivalent to the following equations:

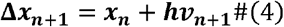

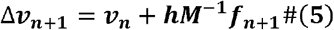

The non-linear force term can be approximated using a Taylor series. The linearization is achieved by replacing the nonlinear force term with its first-order Taylor series approximation, as shown in equation 6:

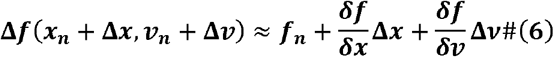

There is a trade-off made to use Taylor series approximation for an adequately close solution because there are indeed 2nd or higher-order time marching methods that can provide more accurate approximation but require more computation time. Then we can get the velocity update as shown in equation 7 by substituting Δ*x* = *h*(*v*_*n*_+Δ*v*)

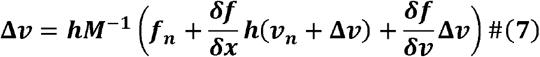

This results in a linear system as shown in equation:

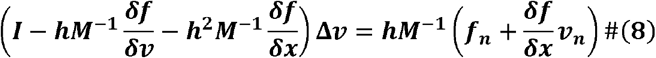

Where *I* ∈ *R* ^3*N*×3*N*^ is the identity matrix, 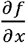 and 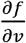 are needed for the internal forces. Internal force derivative with respect to position and velocity is required to be calculated in implicit integration.

Assume *x*_*i j*_= *x*_*i*_−*x*_*j*_ is a normalised vector. The force *f*_*x*_ exerted on a vertex i from a spring connecting particles i and j can be shown in equation 9:

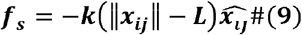

The derivative of force with direction is shown in equations 10:

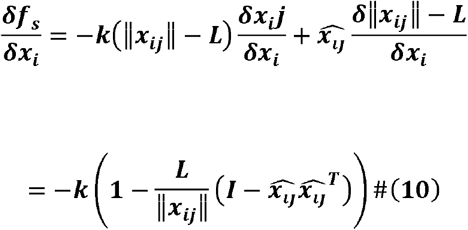

Also, the Jacobian 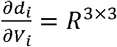 of the damping force is shown in equation 11:

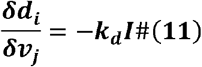

Using these equations, the net Jacobians can be obtained by summing up all the elements yielding 3N x 3N sized matrices. The Conjugate Gradient (CG) algorithm takes a symmetric positive semi-definite matrix A, a symmetric positive definite preconditioning matrix P of the same dimension as A, a vector b and iteratively solves *A*Δ*v*= *B*. The conjugate gradient method used can be described as:

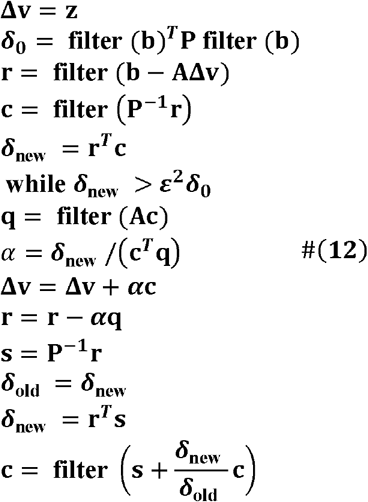

Steps 5 and 15 in algorithm 3.14 maintain the invariant by filtering c before adding it to Δv. The stopping criterion is based on Δb^*T*^ Pb in line 3. The vector r measures the solution error b − AΔv, and should not include error due to the constraints; hence filtering is added in steps 4 and 8. This conjugate gradient method was first proposed in work by Baraff and Witkin [23] and implemented in Visual Studio 2020 and Blender.

#### 2.3.2. Determining system parameters for model

To accurately model the behavior of balloons, stents, and vessels under specific force conditions, the system parameter (k) is calculated in a manner that mimics the Finite Element Analysis (FEA) approach, as established in previous studies [24,25]. The equation can be shown below:

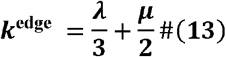

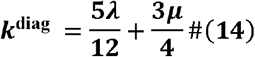

With Lamé constants:

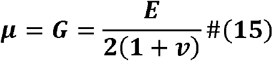

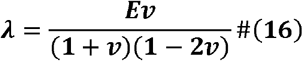

The damping coefficient is determined based on the work by [26,27]. Consider a simple mass-spring-damper system consisting of two nodes with masses *m*_1_ and *m*_2_, where the total mass is denoted as *m* = *m*_1_ + *m*_1_. These nodes are connected by a spring with a stiffness constant k and a rest length. When the mass-spring-damper system is subject to external forces, the damping ratio ζ can be used to analyse the complex oscillations occurring within the springs [28] Critical damping is employed in the fast stent deployment system to prevent unnecessary oscillations and stabilize the system as quickly as possible [24].

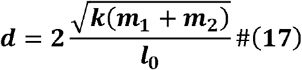

The mass (m) of each vertex is determined approximately by dividing the total mass of objects by the total number of vertices. System parameters such as m, k and damping coefficients can be assigned to the VVI framework developed in Visual Studio 2020 (Microsoft, Albuquerque, New Mexico, USA) in C++. Blender (Blender Foundation, Amsterdam, The Netherlands) modules are then used for the rendering of simulation results and output model as STL (Stereolithography) file.

#### 2.3.3. Collision handling

There are three aspects to collision management: bounding volume hierarchies, fast collision detection, and collision response. The computer graphics (CG) community has developed robust and efficient collision detection algorithms, and in this work, we provide a brief introduction to one such algorithm. More detailed information can be found in the works [25]. Our approach involves representing the 3D collision objects in a hierarchical fashion using bounding volume hierarchies (BVH) to effectively filter out the majority of non-colliding pairs of elements. We then check the distance between each pair of segments in neighboring cells. If the distance *d* between two segments of the discrete surgical thread is smaller than a constant value *d* < *a*, we define that the collision is detected.

Another challenge lies in collision response algorithms, which can be classified into three major categories [29,30]: impulse-based methods [31], constraint-based formulations [32,33] and penalty-based methods [34]. In general, impulse-based methods are more suitable for rigid body collision response due to the requirement of precise dynamic collision detection at each time step. Constraint-based methods result in a more plausible simulation at the cost of additional computation [35,36]. Traditional penalty-based methods, which are the easiest to implement, unfortunately suffer from various issues such as the jitter effect. Tang et al. [29] proposed a novel continuous penalty force model for rigid body collisions. Wang et al. [37] improved upon the penalty method by introducing a continuous penalty averaged force model for real-time application in soft objects. We adapt this approach in our VCI framework. The continuous penalty force created by the collision during the time interval Δ*t* is:

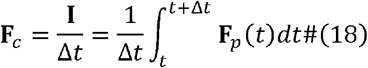

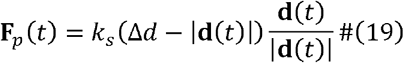

Where *k*_*s*_ is a stiffness constant, **F**_*p*_(*t*) represent the penalty force created at time *t* and 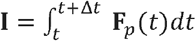 is the impulse produced by the penalty force **F**_*p*_(*t*) during time interval [*t,t*+ Δ*t*]. The continuous penalty force can essentially be taken as the average of the penalty force during a time interval.

#### 2.3.4. Balloon expansion

The expansion of a virtual folded balloon is achieved by applying realistic internal pressure perpendicular to the surface of the balloon, with the pressure distributed to each vertex of the balloon. Additionally, virtual folded balloons based on manufacturers’ guidance and a cylindrical structure with the size of crimped stents were created and discretized with a quadrangular mesh (Fig. 2(d)), then stored for usage.

The equilibrium state of the balloon is achieved by balancing the internal force and the counterforce generated by mass links at each vertex of the quadrangular mesh. The accuracy of balloon expansion is ensured by an expansion test that compares the balloon diameter in the equilibrium state to the balloon manufacturers’ guidance. Regarding the stent, the cylindrical structure represents the crimped stent in its initial configuration, and the simulation of stent implantation is performed by attaching the stent to a balloon of suitable size. Although the structure of the stent can be mapped onto the cylindrical surface, it does not affect the simulation results, and detailed flow analysis in the stented vessel is not included within the scope of this manuscript. For visualization purposes, a stent was manually created as shown in Fig. 2(b).

The simulation of balloon angioplasty and stent implementation was conducted within the VCI framework, which incorporates clinical settings during the PCI procedure, such as balloon size, balloon location, balloon internal pressure, stent size, and stent position. The centerline of the coronary vessel was extracted and imported into the VCI graphical user interface. The operator can maneuver the devices to the region of interest, as depicted in Fig. 2(c). Within the VCI graphical user interface, the operator selects and specifies the desired length and diameter of the stent and balloon to be deployed, as shown in Supplemental Figure 3.

### 2.4. Computational fluid dynamic settings and hemodynamic studies

The coronary artery models, including pre-PCI CTA-reconstructed models, post-PCI CTA-reconstructed models, and corresponding VCI-simulated models, were meshed using CFD-VisCART (ESI Group, Paris, France), employing a stair-step conforming unstructured mesh of the Omnitree Cartesian tree type. Blood flow, assumed to be incompressible, was modeled using the incompressible 3D Navier–Stokes equations solved via the finite volume approach in CFD-ACE+ (ESI Group, Paris, France), with a central differencing scheme for spatial interpolations. The SIMPLE Consistent (SIMPLEC) pressure correction method [39,40] and an algebraic multigrid method for convergence acceleration [41] were used. The flow was assumed to be laminar and blood was modelled as a homogenous and Newtonian fluid with its density and dynamic viscosity of 1060 kg/m3 and 0.004 Pa·s, respectively. For all cases (Pre-PCI CTA-reconstructed models, post-PCI CTA-reconstructed models, and corresponding VCI-simulated models), the vessel wall was approximated as rigid, with non-slip boundary conditions applied. Cardiac-induced wall motion was not incorporated.

In the VCI framework, steady-state BCs are used for the calculation of FFR_CT_. This is based on two reasons. First, the calculation of invasive FFR is based on time-averaged pressure measured over several cardiac cycles. Second, using steady-state BC reduce the computational cost, without losing the predictability of FFR as shown in studies [42,43]. The model of calculating FFR_CT_ is based on left ventricular mass (LVM) based mode. The LVM model quantifies total hyperemic coronary blood flow based on LVM and distributes outlet coronary blood flow based on outlet diameter. Based on the LVM measured from the workstation (CoronaryDoc, Shukun Technology, Beijing, China), the resting state flow Qrest in the right and left coronary branches is calculated as:

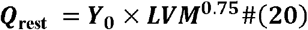

Where Y_0_ is a normalisation constant of 0.71 [44]. The coronary blood flow value in the rest state (Qrest) is assumed to be 16% [42] of that in the hyperemic state. Qhyp can be calculated as:

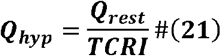

TCRI is the hyperemic factor, and TCRI is 0.16 [43]. Murray’s law between vessel radius is applied. The flow in the coronary branch at ith outlet 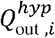 can be calculated according to the following equation:

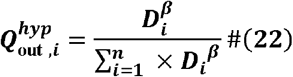

Where *β* is 2.55, *D*_*i*_ is the diameter of each branch outlet, and n is the number of branch outlets [45]. The coronary outlet pressure is set to MAP (MAP=0.4*(SBP-DBP)+DBP) [46] where SBP and DBP are brachial systolic and diastolic blood pressure provided by CoronaryDoc, Shukun Technology, Beijing, China. The coronary artery resistance 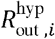 can be calculated as:

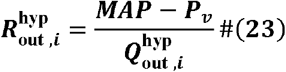

Where 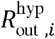 is the resistance of ith outlets, *P*_*v*_ is assume to be 5 mmHg [1]. The same boundary conditions were assigned for each patient case (post-PCI CTA-reconstructed models and the corresponding VCI-simulated models) at the inlet and outlets. Convergence criteria of absolute and relative residual reductions of 1×10^−8^ and 1×10^−5^ were imposed respectively, with convergence typically reached in fewer than 100 iterations.

### 2.5. Comparison metrics

The post-VCI FFR, post-VCI lumen, and post-VCI curvature were compared to the post-PCI FFR, post-PCI lumen, and post-PCI curvature segmented from CCTA. The metrics from post-PCI CCTA were used as a reference. The mean lumen diameter, post-FFR and lumen curvature along the stented coronary vessel were used as comparison metrics.

### 2.6. Statistical analysis

Statistical analysis was performed with the statistical package MedCalc® Statistical Software version 20.116 (MedCalc Software Ltd, Ostend, Belgium). Continuous variables were expressed as mean ± standard error of the mean. The main aim was to access the agreement between post-VCI FFR and post-PCI FFR by using the Bland-Altman method, other metrics such as lumen crosse section and curvature in the stented area also accessed using the Bland-Altman method. The metric of accuracy is defined by the mean difference and the precision is defined as the standard deviation (SD) of the mean.

## 3. Results

### 3.1. Efficiency of VCI

All the computations were executed in Visual Studio C++ on one 11th Gen Intel(R) Core (TM) i7-11800H CPU @ 2.30GHz processor with 32GB of RAM. Simulation of a complete PCI process consists of three steps: pre-dilatation (PRE), stent-deployment (DEPLOY), and post-dilatation (POST). Each step involved three stages: deployment-to-contact (Stage 1), contact-to-balance (Stage 2), and balloon-extraction (Stage 3). Fig. 3 shows the simulation for one representative case which includes all pre-dilatation, stent deployment, and post-dilatation steps. It takes a few seconds to compute each step on a single CPU. For the 10 successfully treated cases, the mean computing times for PRE, DEPLOY and POST processes were 7.93 ± 2.52 s, 8.39 ± 2.66 s and 8.39 ± 1.70 s. The mean computational times for Stage-1, Stage-2 and Stage-3 were 8.78 ± 1.02 s, 6.99 ± 1.11 s and 9.15 ± 0.88 s as it can be seen in Fig. 4, and the overall simulation time for virtual stenting was 24.92 ± 1.00 s.

**Figure 3.**
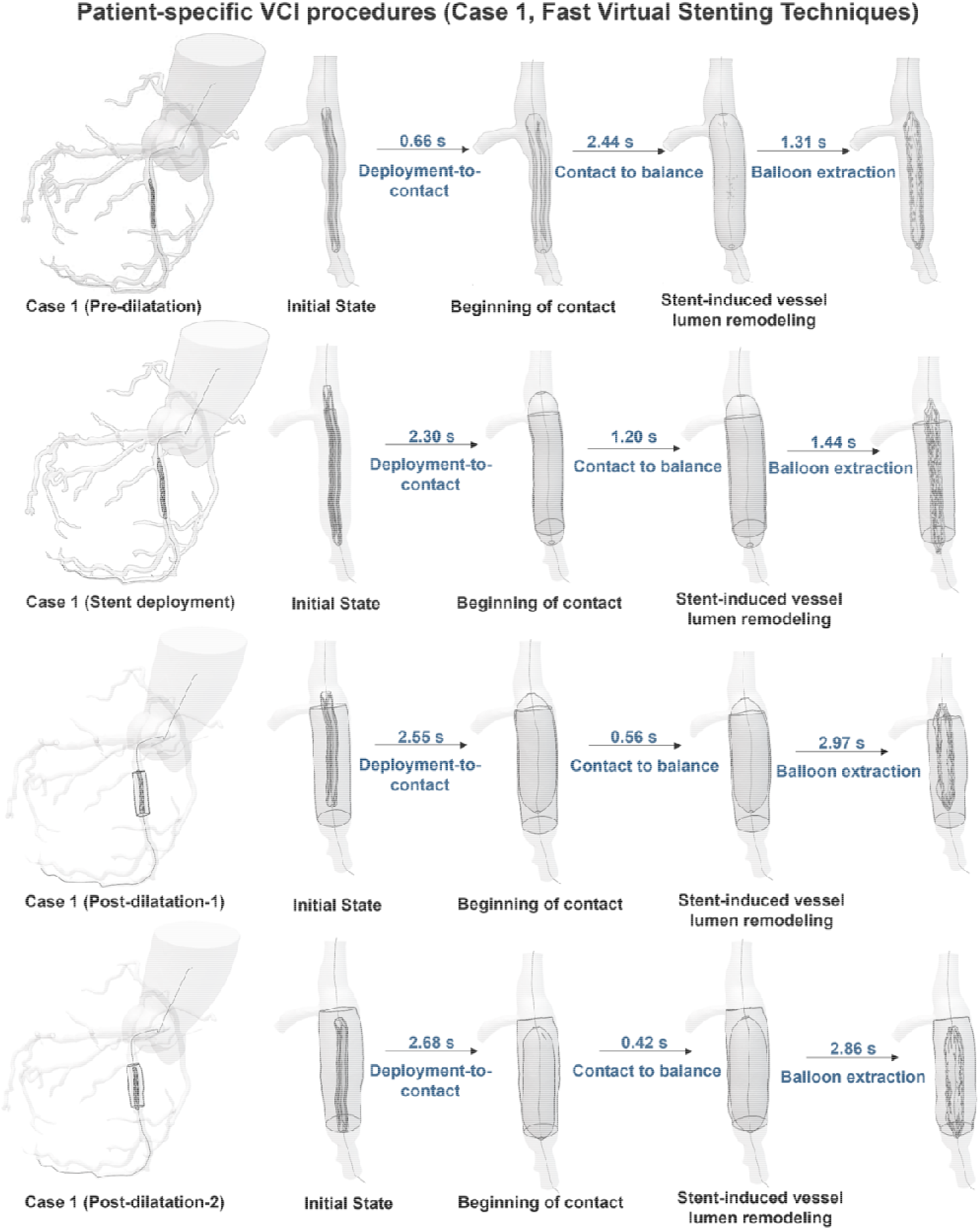
Fast Virtual Stenting Technique for representative case. Four steps are included in the clinical process: pre-dilatation, stent deployment, post-dilatation-1, and post-dilatation-2. We replicated computationally all the steps of the stenting procedure within VCI framework. Three stages are included in each step and the computation time is rapid. For each clinical procedure, there are three main states: initial states, beginning of contact and stent-induced vessel lumen remodeling states.

**Figure 4.**
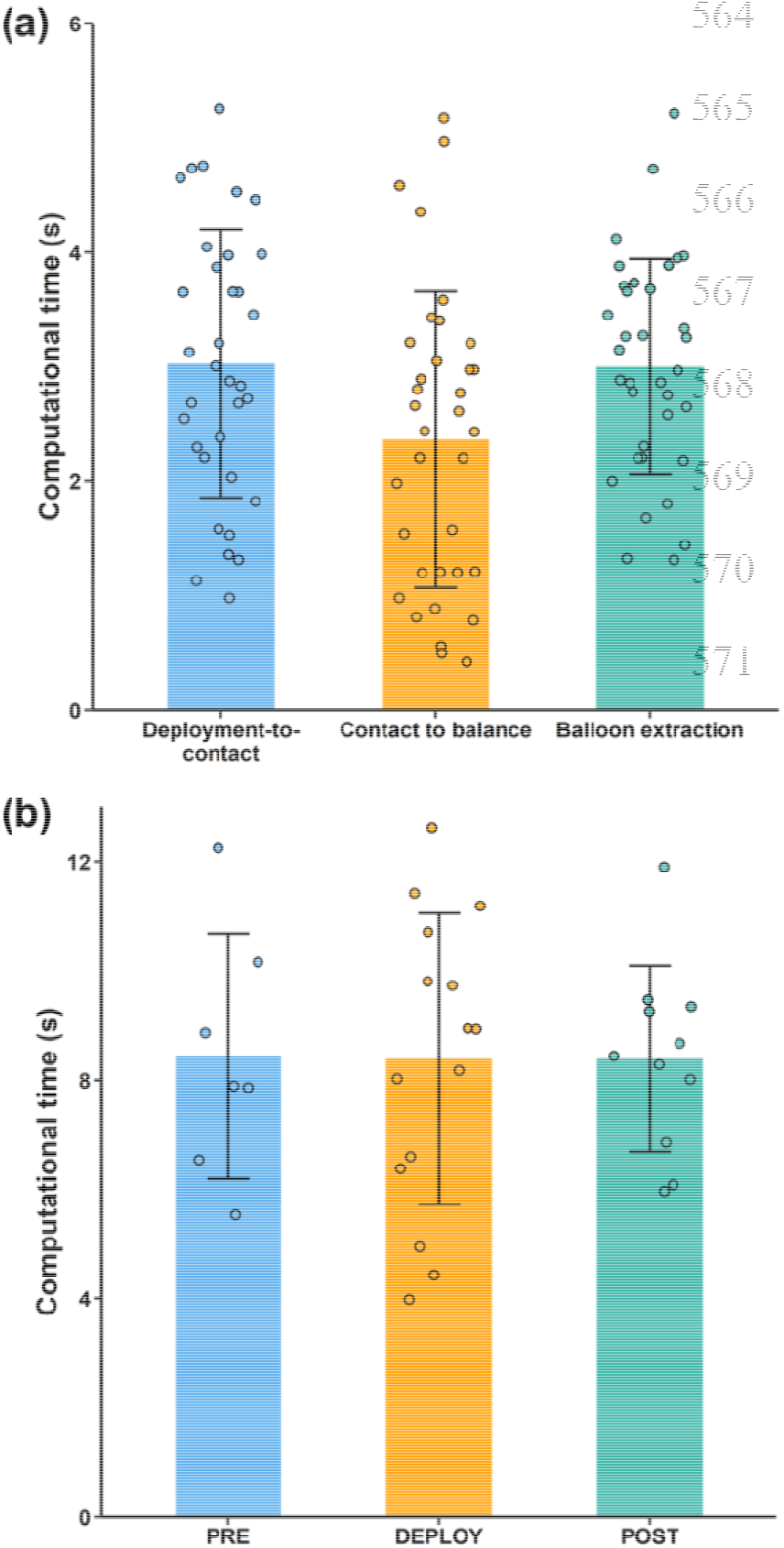
Computational time for fast virtual stenting technique. (a) Computing time for different stages during virtual stenting for all the simulated cases, there are deployment-to-contact, contact-to-balance and balloon extraction stages. (b) Computing time for virtual balloon pre-dilatation (PRE), stent-deployment (DEPLOY), and balloon post-dilatation (POST) for all the simulated cases.

### 3.2. Accuracy of the VCI

Fig. 5 (a) and(b) show both the pre-PCI and post-VCI LAD vessel, lesion vessel cross-section view and calculated FFR based on CCTA reconstruction. The accuracy of VCI was accessed mainly based on FFR_CT by comparing post-VCI FFR to the real post-PI FFR CTA-reference models (CTA model) as illustrated in Fig. 5 (c) and (d). The calculated post-PCI FFR was 0.92 ± 0.09 and the predicted post-VCI FFR was 0.90 ± 0.08 (mean difference: -0.02 ± 0.05 FFR unit; limits of agreement: -0.08 to 0.05) (Fig. 5(e) and (f)). As for the morphological analysis between VCI and the reference model, the CSA and luminal curvature in the stented region are compared (Fig. 6(a)). In the stent deployed region, the centreline along with perpendicular slices were extracted (Fig. 6(b)). CSAs (Area _n_, n = 1 - N) (N was in the scope of 60-100, due to the different lengths of the stented vessel) were compared between the VCI simulation and CTA-reference outcomes, as:

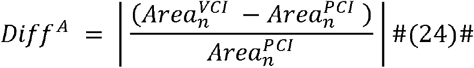

Besides, luminal curvature of the n^th^ points along the centreline (Cn, n=1-N) was evaluated by computing:

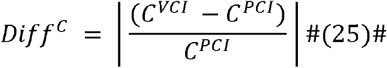

Fig. 6(c) shows comparison of extracted cross-sectional area (CSA) from post-PCI model and post-VCI model in one representative case. The predicted post-VCI CSA is 16.356 ± 4.409 mm^2^ and post-PCI CSA is 17.91 ± 4.84 mm^2^ (mean difference: -1.55 ± 1.89 mm^2^; limits of agreement: -5.22 to 2.12) (Fig. 6(d)), the predicted centreline curvature of stented region is 0.15 ± 0.04 mm^-1^ and post centreline curvature is 0.17 ± 0.03 mm^-1^ (mean difference: -0.02 ± 0.06 mm^-1^; limits of agreement: -0.12 to 0.09) (Fig. 6(e)). The accuracy of the VCI was stratified according to the morphological parameter as well as hemodynamic parameter.

**Figure 5.**
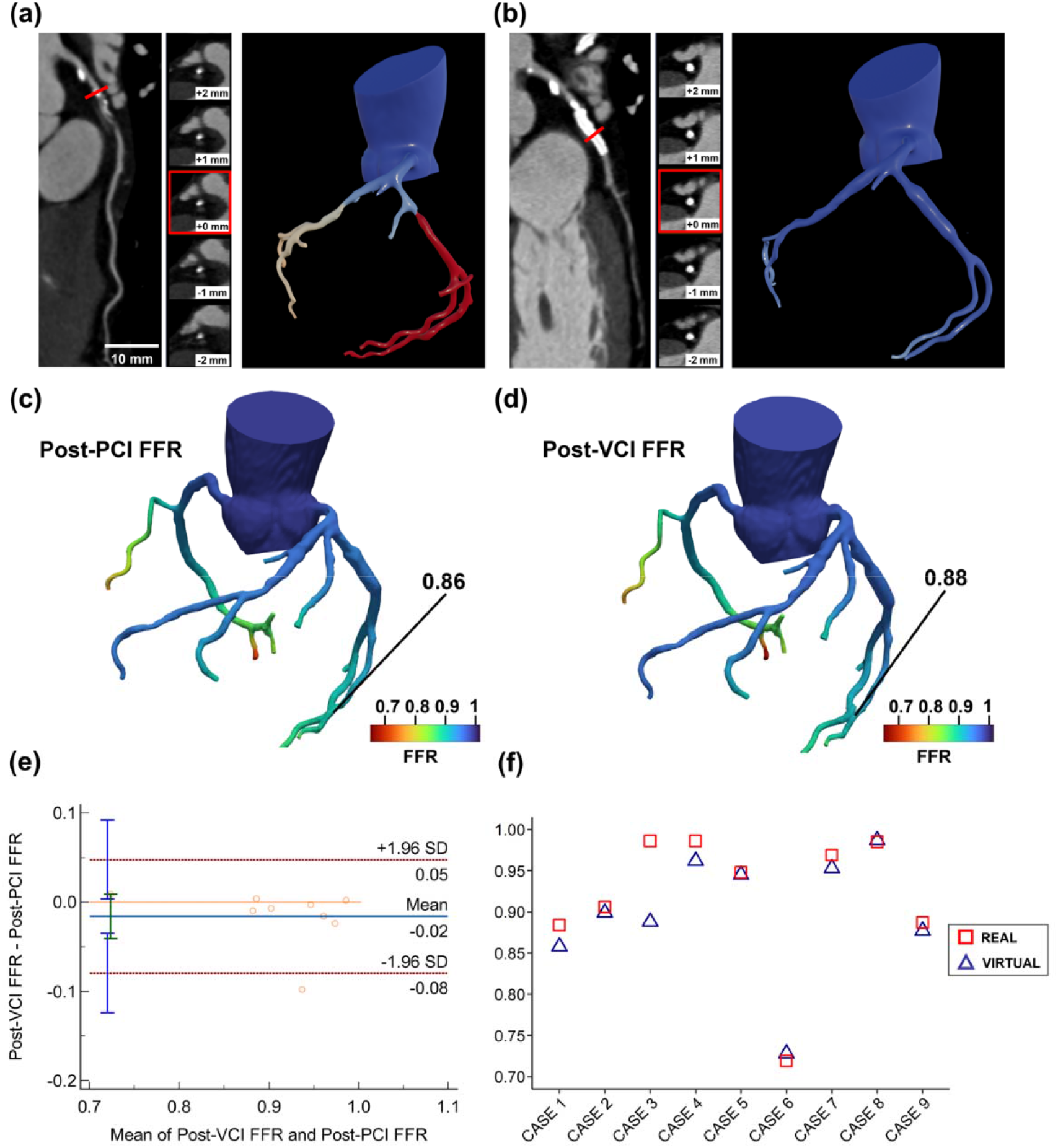
Validation of VCI framework with regard to post-PCI FFR prediction. (a) and (b) left anterior descending (LAD) artery and LAD artery cross section view image along with calculated FFR based on CCTA for pre and post PCI FFR. (c) and (d) Post-VCI FFR and Post PCI FFR, the predicted post-VCI FFR is 0.86 at distal LAD and reference Post PCI FFR is 0.88 at same probe location for one representative case. (e) and (f) Statistical results for the comparison between Post-VCI FFR and Post PCI FFR.

**Figure 6.**
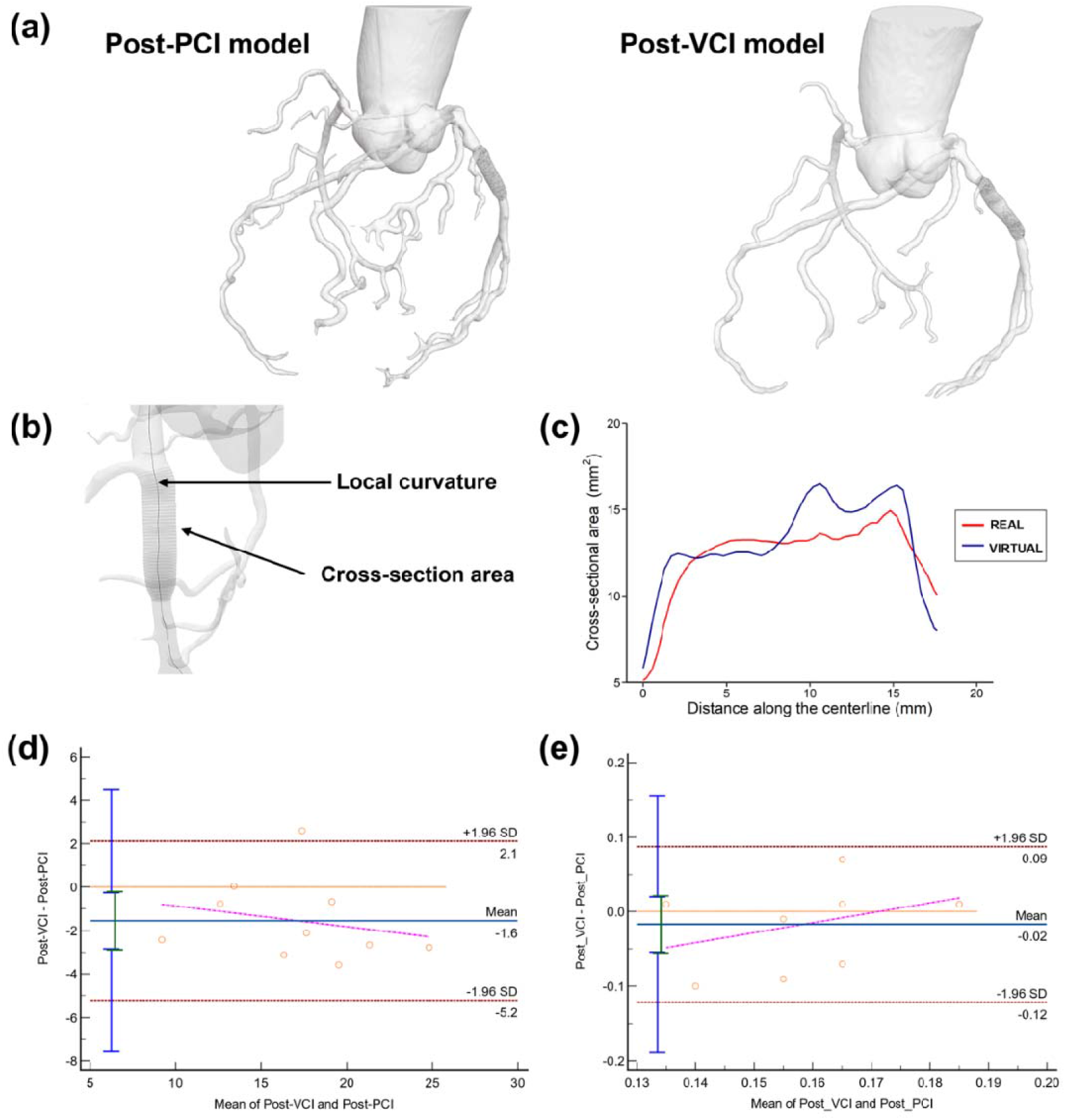
Validation of VCI framework for curvature and crosse section prediction. (a) Crosse section analysis region from post-PCI model and post-VCI model. (b) local curvature and crosse section slicers were extracted. (c) The line graph that shows comparison of extracted cross-sectional area (CSA) from post-PCI model and post-VCI model in one representative case. (d) and (E) Statistical results for both CSA and curvature among all simulated cases.

## 4. Discussion

In this study, we proposed and validated a patient-specific novel VCI technique. T The main results suggest the proposed VCI technique can predict post-PCI FFR with high accuracy. The predicted post-VCI FFR was close to the post-PCI FFR derived from CCTA after PCI. Moreover, the evaluation of morphological parameters (CSA and centreline curvature) demonstrated the rapid VCI prediction is consistent with actual PCI results. In this study, a mean of 24.91 ± 1.00 s was used on a single processor for the fast coronary stenting simulation, which in most cases (n = 8) consists of three steps (PRE, DEPLOY and POST). Previous 3D computational stent-induced coronary vessel remodelling studies were mainly based on the FEM for structural analysis which is not suitable for clinical use due to its high computation requirement [47–49].

Although the fast VCI simulated by using a cubic spline to adjust the cross-sectional diameters to smooth the vessel trajectory based on the size (diameter and length) of the stent proposed by [50,51], this radius correction virtual stenting method can achieve rapid computation of stenting process, the accuracy of such methods might not be insured for the accurate anatomical assessment. The radius correction virtual stenting method might not be ensured for cases of severely calcified vessels, bifurcation stenosis, or aortic-ostial lesions, severe vessel tortuosity, previous revascularization, and in patients with atrial fibrillation, etc [1]. It is important to note that, for patients with pre-PCI FFR less than 0.80, above mentioned scenario occurs frequently. On the contrary, the VCI method can provide a more realistic prediction of morphological outcomes which has the potential to address the above-mentioned challenges.

A more realistic VCI like the one developed in this study that can simulate multiple stent deployment and can adapt to various vessel conditions is therefore required to address the mentioned challenges. Compared to previous coronary stenting algorithms [47–51], the VCI developed in this work has the following advantages:

### 1. The ability to realise complex PCI simulations with rapid computation

As can be seen in Fig 7 two stents were deployed to the bifurcation site known as the culotte technique. In this case, a stent is first developed to the main branch (MV), and the first proximal optimisation technique is applied to the 1st stent to ensure the full expansion of the 1st stent, the 2nd stent is delivered and deployed in the side branch (SB) before side branch structure opening process. The second proximal optimisation technique is used to ensure the contact of 2nd stent with MV. Finally, kissing balloon inflation is performed to avoid the obstruction of the bifurcation vessel. This benchcase shows the potential of VIC to perform complete PCI procedures such as the culotte technique.

**Figure 7.**
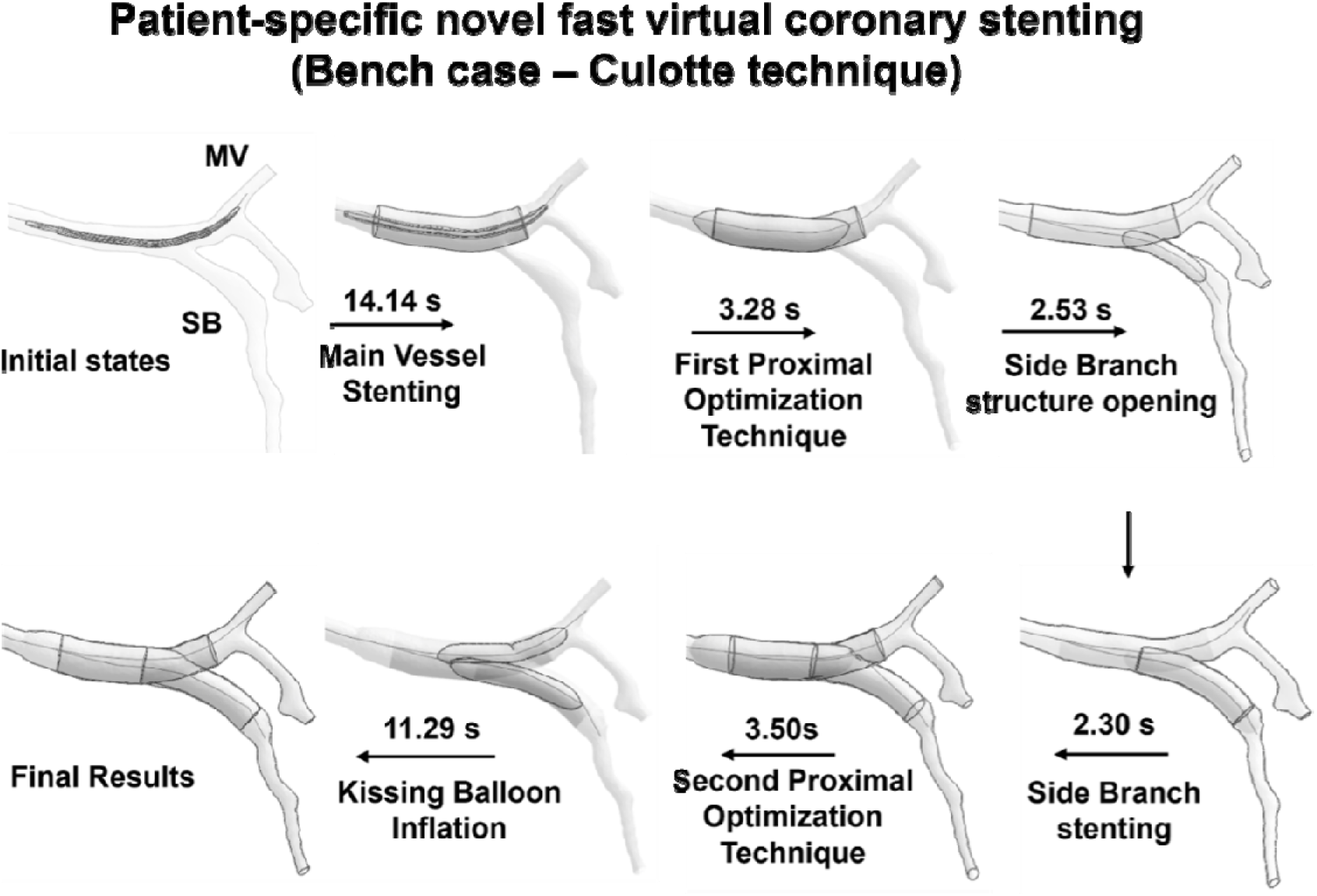
A patient-specific coronary bifurcation geometry was stented with the two stents using Culotte technique.

### 2. The ability to simulate pre-dilatation and post-dilatation in real-time with balloon expansion

Pre and post-dilatation process was simulated in the actual PCI among 8 cases and simulated within the VCI framework proposed. Most of the previous fast VCIs neglected pre-dilatation and post-dilatation which can affect the accuracy of predicted result as it can be seen in the Fig 8, the predicted Post-RC FFR is 0.86 with only stent implation process. Whereas the predicted Post-VCI FFR from the VCI method that contatin pre-dialation and then stent implantation process is 0.88 which is closer to the Post-PCI FFR – 0.89.

**Figure 8.**
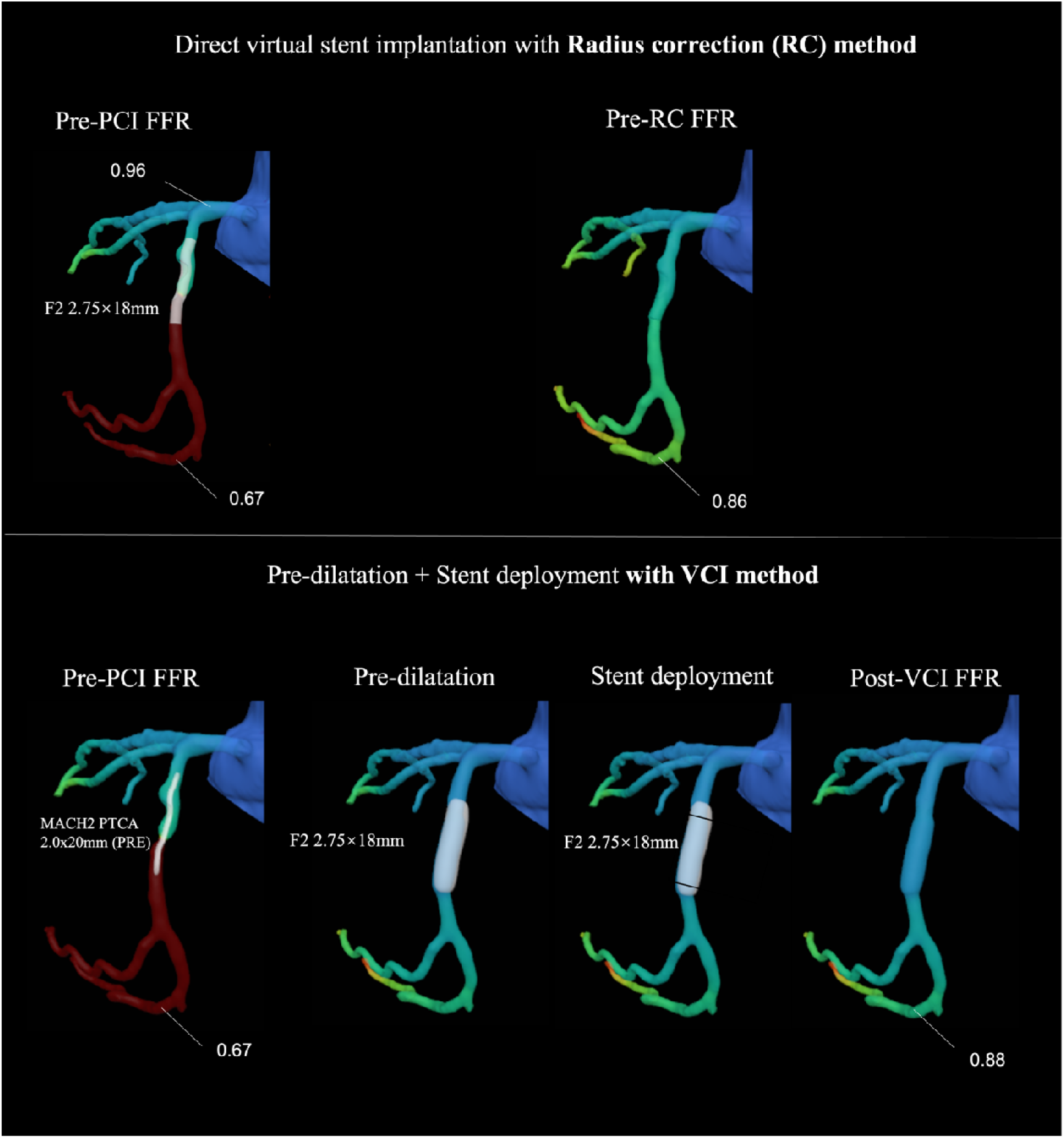
The comparision between direction stent implantation with RC method and VCI method that contain pre-dilatation and stent implantation steps.

### 3. The potential to simulate various vessel conditions

In the previous FE study, the plaque can be simulated by the stress-strain graph based on the constitutive equation with multiple coefficients [14]. A similar idea can be applied to the fast coronary stent deployment algorithm proposed in this study by changing the vessel material parameters, further investigation should be carried out to export the potential of algorithms to mimic different vessel properties. The ability to simulate two-stent techniques in the bifurcation of coronary such as the double kissing crush technique and culotte technique should be examined further.

The study has its own limitations. Perhaps the key limitations are: (1) the number of cases analysed in this proof-of-concept study is modest due to some excluded cases from the automated vessel segmentation process as in rare cases with very tortuous coronary geometries, the segmentation failed. With the future improvement of automated 3D reconstruction algorithms more patients with complex diseases are warranted; (2) the movement of vessels during the cardiac cycle was not considered. (4D simulation) with time make the simulation accurate at the expense of computation time; (3) mechanical contact analysis should be carried out in the future for better modelling of vessel wall response which can replace the stable force in the fast virtual stenting [21]. The material properties of different plaque types should be modelled based on stress–strain curve obtained from the mechanical test; (4) invasive measurement should be used for the validation of FFR_CT and OCT should be used for the validation of plaque segmentation. More validation should be carried out if the relevant data is available; (5) A non-Newtonian flow model should be adopted for the FFR calculation to consider the effect of blood cells which are the main factors behind blood viscosity.

### 5. Summary

Non-invasive physiological assessment (post-PCI FFR_CT_ prediction) and anatomical assessment (detailed coronary stenting simulation) can facilitate pre-procedural planning of PCI by evaluating the degree of functional revascularization and predicting stented coronary vessels. The VCI framework proposed in this work achieved physiological assessment by accurately predicting post-PCI FFR_CT_, and anatomical assessment by simulating detailed stenting procedures in real-time.

The present framework could contribute to functional coronary angiography (FCA) derived from computed angiograms to facilitate non-invasive PCI planning and the present framework has the protentional to assist in improving the efficacy and safety of PCI by performing complex interventions in real time.

## Ethics

This study was approved by the Institutional Review Board of the Chinese PLA General Hospital (S201703601)

## Data accessibility

All data generated or analysed during this study are included in the manuscript and supplementary tables and figures.

## Statements and Declarations

The authors declare no conflict of interest.

## Author contributions

ML.: conceptualisation, data curation, formal analysis, funding acquisition, investigation, methodology, writing original manuscript, reviewing and editing;

CL.: formal analysis, investigation, methodology, reviewing and editing;

XZ.: formal analysis, investigation, methodology, reviewing and editing;

XW.: formal analysis, investigation, methodology, reviewing and editing;

QL.: formal analysis, writing original manuscript, reviewing and editing;

RT.: conceptualisation, formal analysis, investigation, methodology, reviewing and editing;

YV.: conceptualisation, funding acquisition, formal analysis, investigation, supervision, reviewing and editing.

DC.: conceptualisation, funding acquisition, formal analysis, investigation, supervision, reviewing and editing.

## Acknowledgements

The authors would like to thank Shukun (Beijing) Network Technology Co. Ltd. for providing additional computation resources and technical support.

