## Supplementary Material for "A Novel Computational Pre-Procedural Planning Model for Coronary Interventions Based on Coronary CT Angiography"

**Supplemental Figure 1.** Study design flow chart

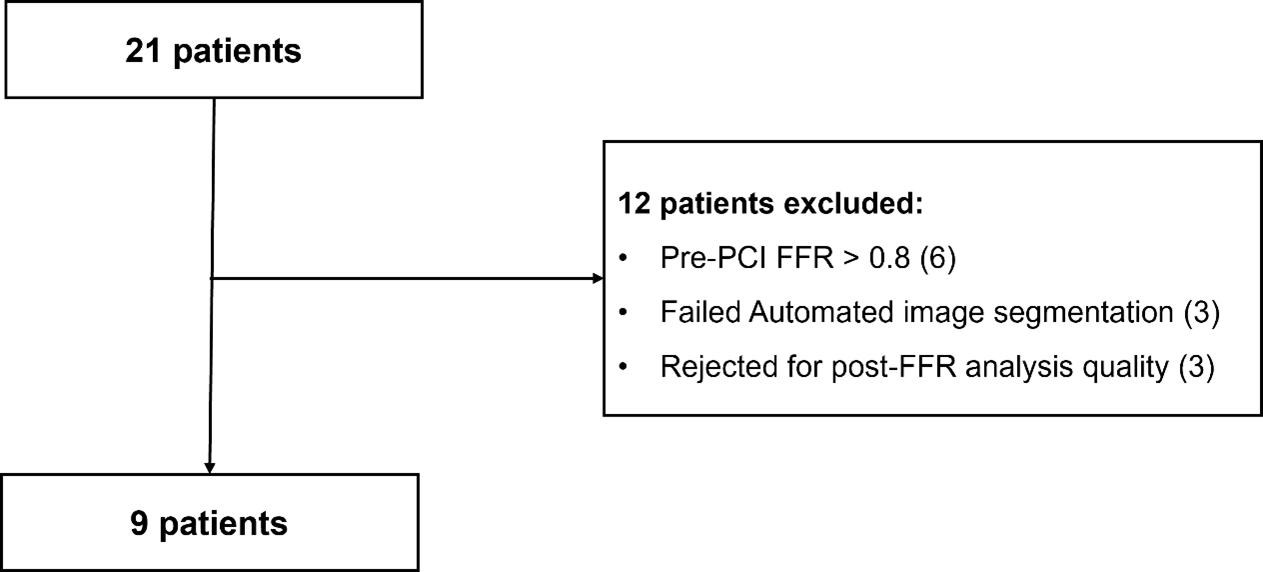

**Supplemental Table 1*.*** Clinical PCI procedures for 9 cases

| **Steps** | **Case 1** | **Case 2** | **Case 3** | **Case 4** | **Case 5** |
| --- | --- | --- | --- | --- | --- |
| 1 | LCX pre-dilatation MACH2 PTCA 2.0×20mm @ 10 atm | LAD pre-dilatation SPRINTER 2.0×12 @ 12-14 atm | LAD pre-dilatation MicroPort Pioneer 2.5 × 15 mm @ 14 atm | LAD pre-dilatation Pioneer 2.5 × 15mm @ 8-10atm | LAD pre-dilatation Pioneer 2.5×15 mm @ 8-10atm |
| 2 | LCX stenting F2 2.75×18mm @ 10-16 atm | LAD stenting Firebird 3. 5×29mm @ 10-16 atm | LAD stenting JWMS EXcel 3.5 × 14 mm @ 12 atm | LAD stenting Firebird2 3.0×33mm @ 8atm | LAD stenting Firehawk 2.75× 29 mm @ 8atm |
| 3 | LCX post-dilatation GOODMAN2 5×13mm @ 12 atm | LCX Post-dilatation NC SPRINTER 3.5×12mm @ 14-18atm |  | LC× Post-dilatation NC SPRINTER 3.0×15mm @ 16-18 atm | LAD stenting NaNo 3.0 mm × 24mm @ 8atm |
| 4 | LCX post-dilatation NC Sapphire 2.75×12mm @ 16-20atm |  |  |  | LC× Post-dilatation NC SPRINTER 3.0 × 15mm @ 16-18 atm |
|  | **Case 6** | **Case 7** | **Case 8** | **Case 9** |  |
| **1** | LCX pre-dilatation Pioneer 2.0×15mm @ 8-10atm | dRCA pre-dilatation Goodman Lacrosse 2.5×15mm @ 10 atm | RCA pre-dilatation NINI TREK 2.0×15mm @ 10-12atm | RCA pre-dilatation Sequent 2.0×15mm @ 10-12atm |  |
| **2** | LCX stenting Boston Scientific Promus PRENIERTM 2.25×20mm 8 atm | dRCA stenting microPort Firebird2 2.75×13mm @ 12 atm | RCA stenting XTEXCE Xpedition 2.5×33mm @ 8atm | RCA stenting Resolute Integrity 3.0×15mm @ 10-12atm |  |
| **3** | LC× stenting Resolute Integrity 2.78×15mm @ 9atm | dRCA_SB stenting microPort Firebird2 3.5×18mm @ 12 atm | RCA Post-dilatation NC SPRINTER 2.5×15mm @ 16-20atm | RCA Post-dilatation NC Sprinter3.0×9mm @ 16-20atm |  |
| **4** | LCX Post-dilatation NC SPRINTER 2.5× 12mm @ 12-16atm | dRCA Post-dilatation Goodman Lacrossc Powered 2.75×10mm @ 10 atm |  |  |  |
| **5** | LCX Post-dilatation NC SPRINTER 2.75×12mm @ 12atm-16atm | dRCA_SB Post-dilatation Goodman Lacrosse Powcred 3.5×10mm @ 10 atm |  |  |  |
| **6** | R pre-dilatation Pioneer 2.0×15mm @ 8-10 atm |  |  |  |  |
| **7** | R stenting Resolute Integrity 2.25×18mm @ 8 atm |  |  |  |  |
| **8** | R Post-dilatation NC SPRINTER 2.5×12mm |  |  |  |  |

**Supplemental Figure 2 VCI graphical user interface**

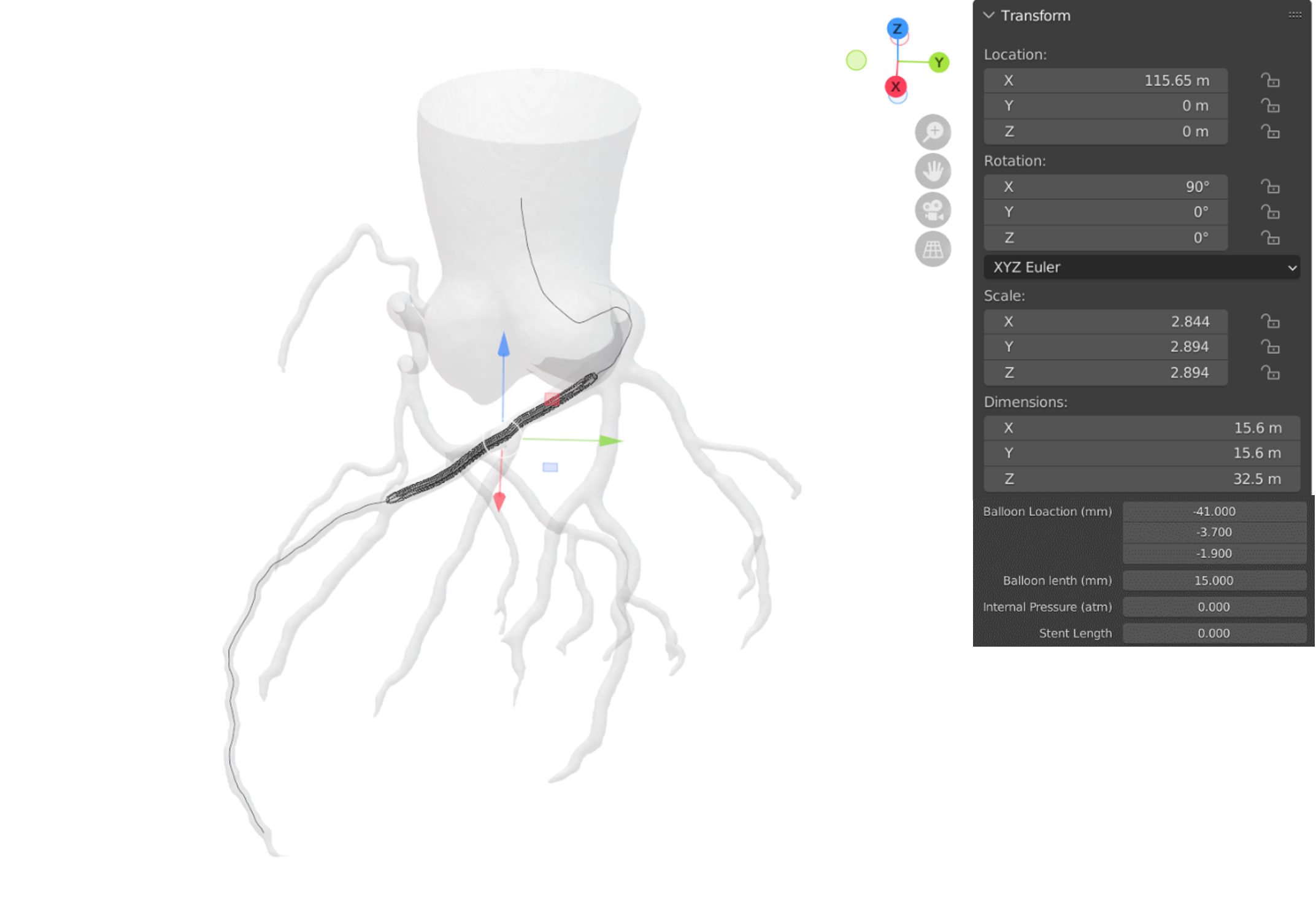

**Supplemental Figure 3** Overview of three virtual fast pre-operative processes (A, B, C). (A) Virtual folding process, the reconstructed balloon catheter is pressed into a star-like shape with three wings. (B) Virtual pleating process, the wings are folded over. (C) Virtual crimping process, the stent is created to fit to the pleated balloon catheter (Blender render module used to produce realistic visual effect).

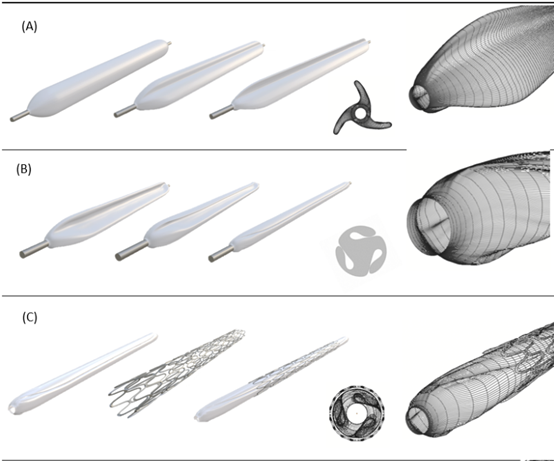
